# Two Key Actinomycetota Taxa in the Human Gut Microbiota are Associated with *Schistosoma mansoni* Infection Burden

**DOI:** 10.1101/2024.12.03.626529

**Authors:** Francis Appiah-Twum, Lydia Okyere, Jeffrey Gabriel Sumboh, Dickson Osabutey, Rahmat Bint Yusif Ismail, Hilda Darko, Yvonne Ashong, Michael D. Wilson, Jewelna Akorli

**Affiliations:** Department of Parasitology, Noguchi Memorial Institute for Medical Research, University of Ghana, P.O. Box LG 581, Legon, Accra, Ghana; Department of Pathobiology, University of Illinois at Urbana-Champaign, Urbana, Illinois 61802, United States; School of Tropical Medicine and Global Health, Nagasaki University, 1-14 Bunkyo, Nagasaki City 852-8521 Japan

**Keywords:** *Schistosoma mansoni*, infection burden, acute and chronic schistosomiasis, human gut microbiome, dysbiosis, Actinomycetota, *Bifidobacterium*, *Collinsella*

## Abstract

In this study, we sought to identify key microbial taxa associated with human gut dysbiosis during *S. mansoni* infection and whether the changes are linked to the intensity of helminth infection. Stool samples were obtained from 20 persons infected with schistosomiasis and an equal number of uninfected persons from an endemic rural community in Ghana. Infection intensity was scored as egg count per gram (EPG) using the Kato-Katz method. Positive stool samples were further stratified as low-moderate (<400 EPG, n=15) and high (>400 EPG, n=5) infection burden. The composition and diversity of the gut microbiota and potential microbial markers associated with *S. mansoni* infection intensity were determined from 16S rRNA amplicon sequence analyses. No difference in ß-diversity was observed between positives and negatives (PERMANOVA: R^2^= 0.012, *p*= 0.723), although there was an increased abundance of *Bifidobacterium* (*p*= 0.008) in infected stool samples compared to the negatives. Further analyses showed that *Bifidobacterium* (*p*= 0.003) and *Collinsella* (*p*= 0.029) were elevated considerably among the low-moderate infected samples, while the pathobiont *Escherichia-Shigella* was reduced (*p*= 0.0078). Our findings show that intestinal schistosomiasis results in human gut microbiota dysbiosis, which is only distinguished when the intensity of infection is considered, with two key *Actinomycetota* species assuming importance depending on the infection burden.

**Author Summary:** This study investigates the relationship between Schistosoma mansoni infections, a major cause of intestinal schistosomiasis, and the human gut microbiome. Using samples from an endemic region in Ghana, the research examines how infection intensity impacts gut bacteria. The findings reveal that certain beneficial bacteria, such as Bifidobacterium and Collinsella, become more abundant in cases of low to moderate infection, potentially maintaining immune regulation and gut health. However, these effects are not seen in high-infection instances, possibly due to the aggressive hallmarks of high-intensity helminth infections. Understanding these dynamics could be pivotal for developing microbiome-based interventions to improve treatment outcomes for schistosomiasis and similar parasitic infections. This study sheds light on the complex interplay between infectious parasites and gut microbes, emphasising the promise of microbiome research in enhancing public health efforts in areas where parasitic diseases persist.

## Introduction

Schistosomiasis is a major neglected tropical disease that poses great challenges in terms of control, morbidity and mortality [1]. Infection with Schistosoma causes iron deficiency anaemia, malnutrition, stunted growth, increased susceptibility to other illnesses, and impaired academic performance in school-aged children. [2]. Schistosomiasis remains a significant intestinal infection in many endemic countries even after several years of mass drug administration with praziquantel [3]. In these regions, factors such as inadequate sanitation, low income, limited access to education, the presence of intermediate hosts, suitable climatic conditions, and frequent water-related activities for work and leisure all contribute to persisting transmission of schistosomiasis.

Mature adult *Schistosoma mansoni* (*S. mansoni*) worms reside in the mesenteric veins of their human host, primarily around the colon, where they reproduce and shed eggs. These eggs migrate through the host’s tissues to reach the lumen of the intestines and are excreted into the soil with the passage of faecal matter. Once in freshwater, Schistosoma eggs hatch into miracidia, which invade the intermediate snail host and undergo two generations of sporocyst development to produce cercariae. Upon release from the snail, the infective cercariae swim and penetrate the human host skin, shedding their tails to become schistosomula. Schistosomula migrate through the venous circulation to the lungs and heart. They subsequently mature in the liver and egress into the portal venous system. Upon maturation, male and female adult worms copulate and reside within the mesenteric venules to perpetuate the life cycle. Without treatment, *S. mansoni* infection evolves into a chronic disease of the bowel due to the formation of egg granulomas in the tissue and subsequent fibrosis as eggs become trapped in the hepatic sinusoids and intestinal wall [4].

*S. mansoni* eggs share the same niche with the gut microbiota; as such, their presence directly affects gut microbiome homeostasis [5]. In mouse models, *S. mansoni* infection leads to significant and widespread alterations in the overall structure of the host gut microbiome [6]. These changes coincide with parasite egg deposition within the intestinal mucosa and the migration of parasite eggs through the intestinal lumen [7].

The gut microbiome is crucial for modulating immune responses [8] and plays a significant role in host metabolism, including nutrient absorption and the production of metabolites essential for maintaining intestinal health [9]. Investigating how helminth infections alter the microbiome is important for understanding the underlying mechanisms of disease progression. These microbial shifts may also influence treatment efficacy [10], potentially providing insights into why mass drug administration with anthelmintic has not completely eradicated schistosomiasis in endemic regions.

Alterations in the microbiome composition of taxa linked to immune activation and inflammation influence host immune responses, potentially contributing to the modulation of parasite loads, severity of infection and long-term morbidity in schistosomiasis-infected individuals [11]. Despite increasing attention on the role of the gut microbiome in health and disease, the specific effects of varying parasite burdens on the local microbial ecosystem remain largely unexplored.

To address this gap, this study investigated how different intensities of *S. mansoni* infection impact gut microbiome composition. Our objective was to understand patterns of dysbiosis associated with varying infection burdens, aiming to enhance the understanding of host-parasite-microbiome interactions in schistosomiasis.

## Methods

### Ethical statement

This study was approved by the Noguchi Memorial Institute for Medical Research Institutional Review Board (NIRB ID #: 100/16-17), the National Institute of Allergy and Infectious Diseases Division of Microbiology and Infectious Diseases (NIAID DMID, Protocol approval # 17-0061), and the Council for Scientific and Industrial Research (CSIR) Institutional Review Board (RPN 008/CSIR-IRB/2017). This was a nested case-control study within the Noguchi Memorial Institute Initiative for NTDs Elimination (NIINE) project. Written informed consent was obtained from all participants before the commencement of the study. Children aged 3-17 years were enrolled in the study following written informed consent from their parents or caregivers and assent from the children themselves. This study was conducted in accordance with the Ghana Public Health Act, 2012 (Act 851) and the Data Protection Act, 2012.

### Sample collection and parasitological analyses

Stool samples for this study were collected from participants in Nyive, a rural community in the Volta Region of Ghana (6°45’28.4”N 0°34’47.2” E). *S. mansoni* infection was determined using the Kato-Katz stool preparation technique [12]. Two experienced technologists examined each stool sample independently, and a third blind-checked for quality control and to resolve cases of major discordance in readings between the two microscopists. Both 10x and 40x objectives were used to observe *S. mansoni* eggs. The infection burden was categorised as either light-moderate (<400 EPG) or heavy (>400 EPG) according to the parasite egg count observed in one gram of stool sample. Twenty *S. mansoni*-positive individuals were selected, with an equal number of *S. mansoni*-negative individuals. Samples of participants with coinfections with other intestinal parasites were excluded from this study.

### DNA isolation and 16SrRNA gene sequencing

Genomic DNA was extracted from the selected stool samples using the Qiagen Stool Extraction Kit (Qiagen, Hilden, Germany) according to the manufacturer’s instructions. DNA concentration was measured using a Qubit 3.0 fluorometer (Invitrogen, Carlsbad, CA, USA). The V3-V4 region of the 16S rRNA gene was then amplified by PCR using Phusion® High-Fidelity PCR Master Mix (New England Biolabs Inc., Ipswich, MA, USA). PCR products of approximately 350 base pairs were visualised on a 2% agarose gel stained with SYBR Green and purified using a Qiagen Gel Extraction Kit (Qiagen, Hilden, Germany).

Amplicon libraries were constructed using the NEBNext® UltraTM DNA Library Prep Kit for Illumina, following the 16S Metagenomic Sequencing Library Preparation protocol (Illumina™, Inc., San Diego, CA, United States) [13], and the concentrations were determined using the Qubit 3.0 fluorometer and PCR. Paired-end sequencing of the libraries was performed on the NovaSeq 6000 Illumina platform following standard Illumina sequencing protocols to generate paired-end reads of 250 bp.

### Sequence data analyses

Using Quantitative Insights into Microbial Ecology version (QIIME2) version 2020.8.0 [14], paired-end reads were merged and trimmed to remove low-quality reads to perform taxonomic assignments and diversity analyses. Specifically, diversity within each group (α-diversity) was estimated with indices including Gini-Simpson, Species richness and Shannon entropy in the *microbiomeSeq* R package [15] to determine the effective number of species within infected and uninfected groups [16]. Beta diversity was investigated using permutational multivariate analysis of variance (PERMANOVA) based on weighted unifrac distances in the R *vegan* package [17,18]. Weighted unifrac distances were also used to quantify the compositional dissimilarities between infected and uninfected samples while considering the phylogenetic relatedness of the taxa within infection groups. Non-metric multidimensional scaling (NMDS) was used to visualise the clustering of bacterial species in the infected and uninfected samples [18].

### Microbial Biomarker Discovery

Linear discriminant analysis (LDA) was employed within the *MicrobiotaProcess* package in R (version 4.1.3) [18,19] to establish the association between the abundance of host-microbial taxa and *S. mansoni* infection. A taxon was considered a potential biomarker if it exhibited an LDA score ≥ 3 and a statistically significant association (p ≤ 0.05 for the Wilcoxon rank sum test and p ≤ 0.01 for the Kruskal-Wallis test). A stand-alone Wilcoxon rank sum test was also used to compare differences between microbiota abundance across *S. mansoni* infection intensity groups (i.e. No Infection, Low-Moderate and High). To account for multiple comparisons, the Benjamini-Hochberg correction procedure was applied. A Benjamini-Hochberg adjusted p-value of ≤ 0.05 was considered statistically significant.

## Results

### Microbial diversity between groups based on *S. mansoni* infection

The assessment of alpha diversity revealed a slight increase in the effective number of species in individuals infected with *S. mansoni*. However, this difference was not statistically significant when compared to uninfected samples (p > 0.05) (Fig1A). These results suggest a comparable complexity of sample biodiversity among *S. mansoni*-infected and uninfected samples. Similarly, the beta diversity analysis showed no significant structural or compositional differences between the microbial communities of the infected and uninfected groups (PERMANOVA: R^2^= 0.012, *p*= 0.723). This lack of dissimilarity was further confirmed by NMDS analysis, which produced a highly homogeneous plot with no distinct clusters or patterns in either group (Fig 1B).

**Fig 1:**
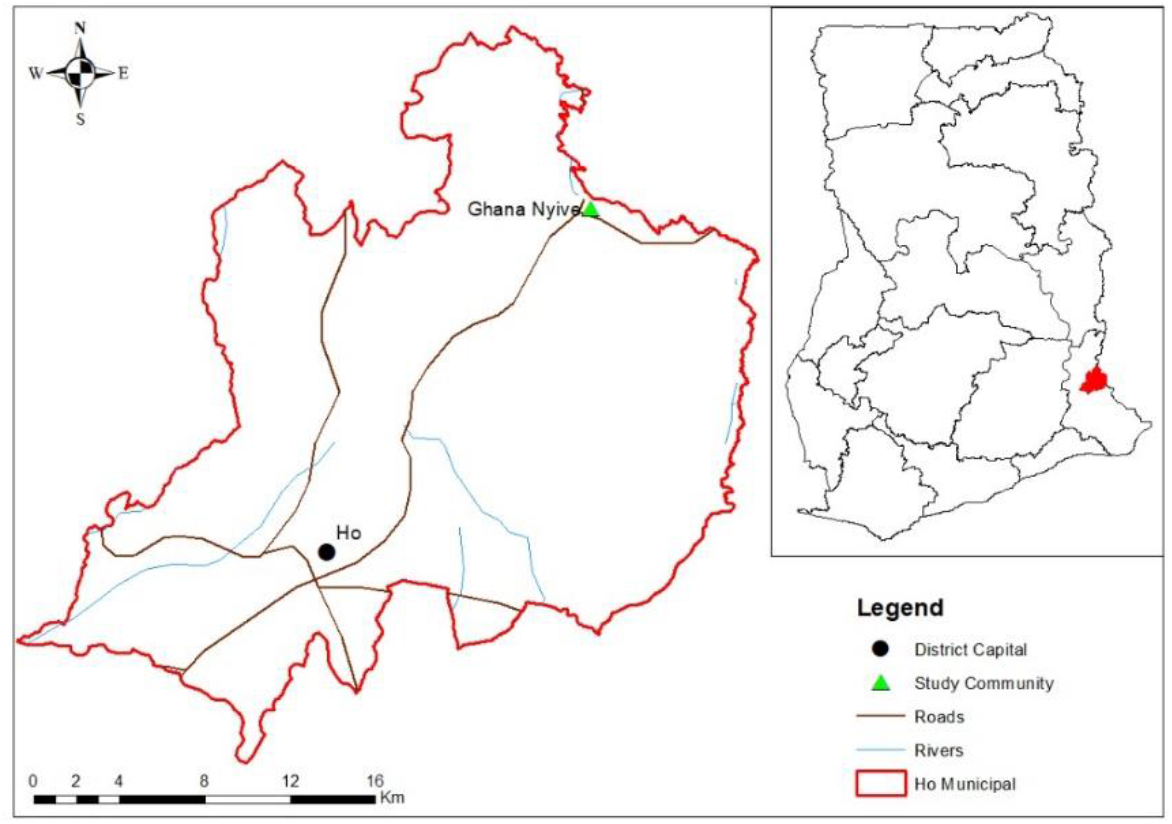
Map of the study site. Nyive. Located in the Ho Municipality, Nyive is a rural community in the Volta Region of Ghana (6°45’28.4”N 0°34’47.2” E).

### Taxonomic identification and microbial biomarker discovery

In the taxonomic characterisation of the gut microbiome, we focused exclusively on taxa exhibiting a relative abundance greater than 1%, as these taxa represent the predominant members of the microbiome. The relative abundances were assessed at multiple taxonomic levels and were subsequently plotted at the phylum and genus levels (Fig 2). Taxonomic composition analysis revealed a similar distribution between infected and uninfected samples at the phylum level (*p>0*.*05*). Analysis of all samples showed that Bacillota (formerly Firmicutes) (68.89%), Actinomycetota (formerly Actinobacteriota) (13.51%), Verrucomicrobiota (9.39%), Bacteroidota (5.22%), and Pseudomonadota (formerly Proteobacteria) (3.00%) were the most abundant phyla across the sample groups. Collectively, these 5 phyla included 43 genera, with the majority (28 out of 43) classified under Bacillota. The phylum Actinomycetota included 6 genera, including *Bifidobacterium* and *Collinsella*, while the remaining phyla contributed to fewer than 2 genera. Notably, 21 of the 28 genera within Bacillota belonged to the Clostridia class, which predominantly exhibited higher relative abundances in samples infected with *Schistosoma mansoni*.

**Fig 2:**
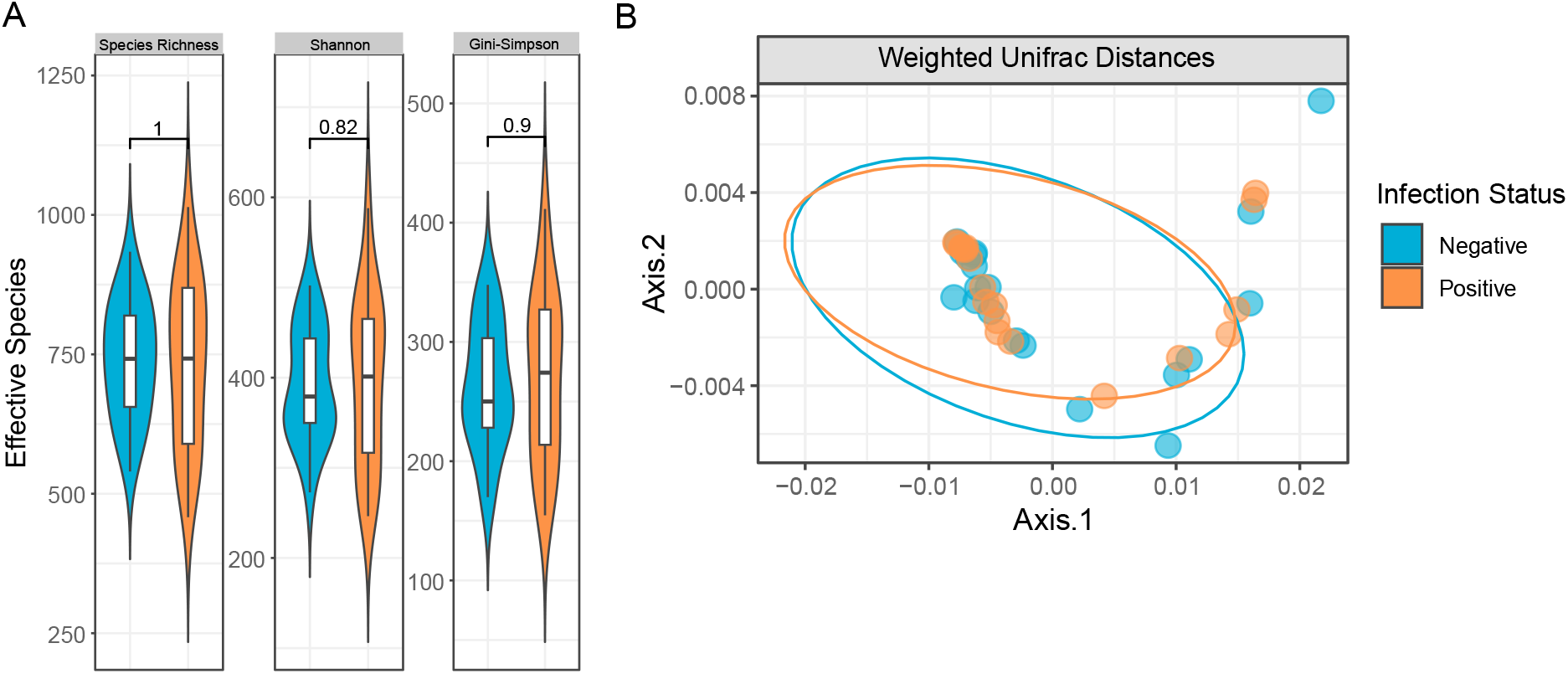
Microbial diversity of study participants based on *S. mansoni* infection status. Alpha diversity analysis using Species richness, Shannon entropy and Gini-Simpson diversity indices **(A)**. Violin plots representing the alpha diversity of *S. mansoni* infected and uninfected individuals. Shannon and Gini-Simpson indices were calculated to represent the effective number of species. **(B)**. Non-metric multidimensional scaling (NMDS) plot of microbiome structure in *S. mansoni* infected and uninfected individuals based on Weighted unifrac distances. The *S. mansoni* infected group is represented in orange, while the uninfected group is represented in blue.

Linear discriminant analysis (LDA) was employed to identify taxa contributing to differences in relative abundance between *Schistosoma mansoni*-infected and uninfected samples observed during taxonomic characterisation. The study revealed a significant increase in the abundance of *Bifidobacterium* (*p*= 0.008) and *Succinivibrio* (*p*= 0.007) in the infected group (LDA score > 3). Although the overall increase in relative abundance across sample groups was modest, the effect size was substantial. This suggests that despite their low abundance, these taxa may play a crucial role in driving the compositional shifts in the microbiota of infected individuals (Fig. 3).

**Fig 3:**
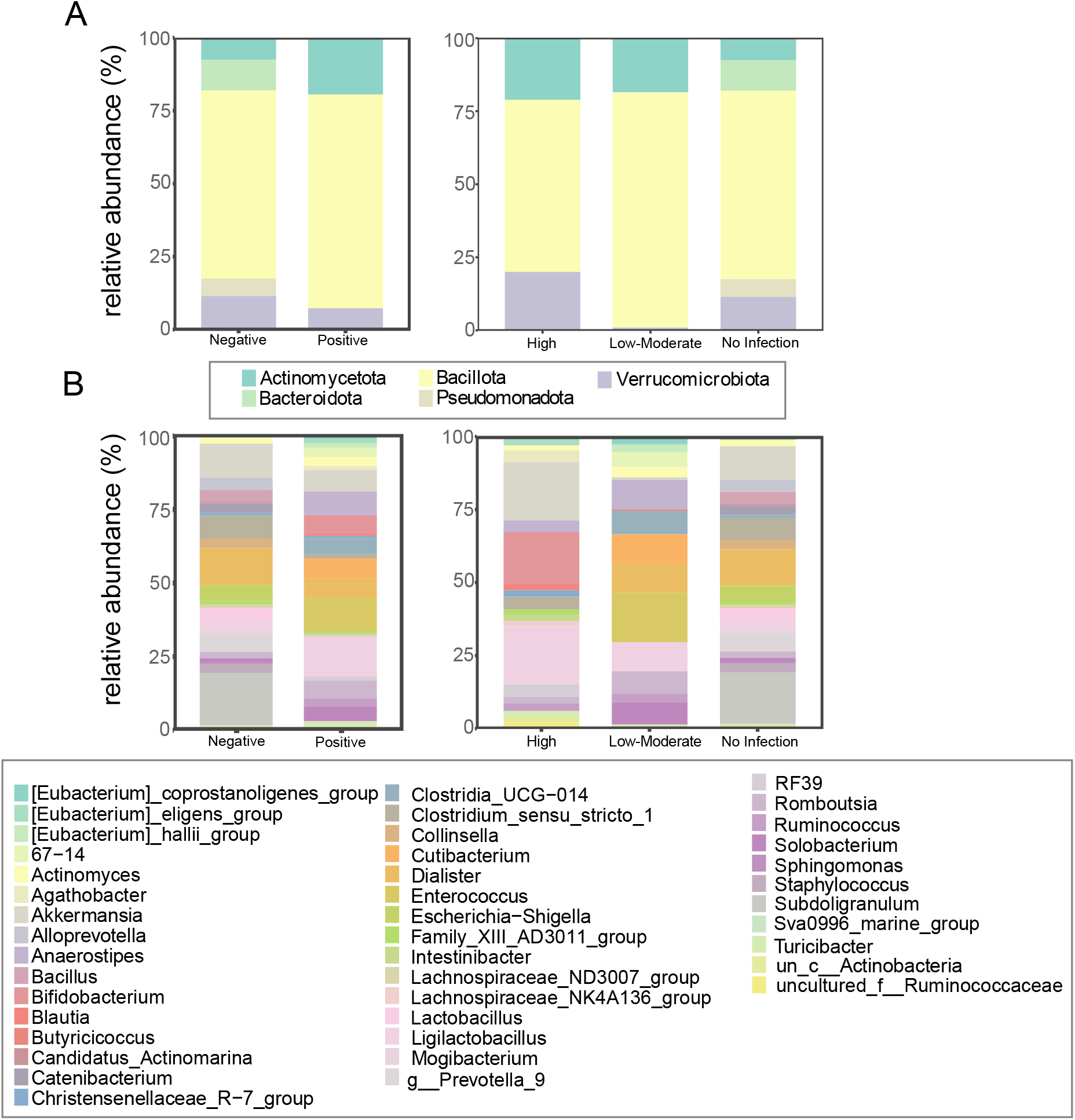
Comparison of the distribution of highly abundant taxa according to *S. mansoni* infection status and infection intensity at the (A). phylum level, and (B). Genus level. Samples with relative abundances greater than 0.01 were classified as highly abundant taxa.

**Fig 4:**
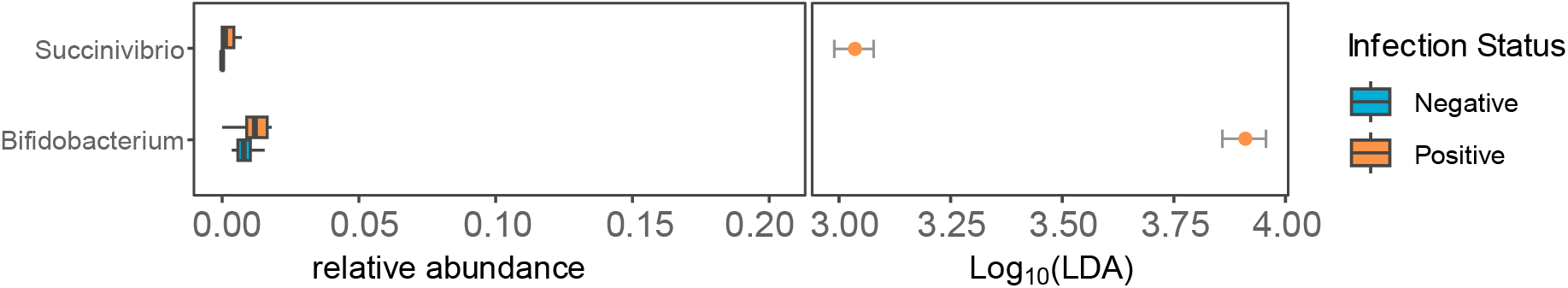
Linear discriminant analysis (LDA) for bacteria genera between *S. mansoni*-infected and uninfected individuals. Plot outlining significantly associated microbial taxa within *S. mansoni* infected (orange) and uninfected (blue) samples.

**Fig 5:**
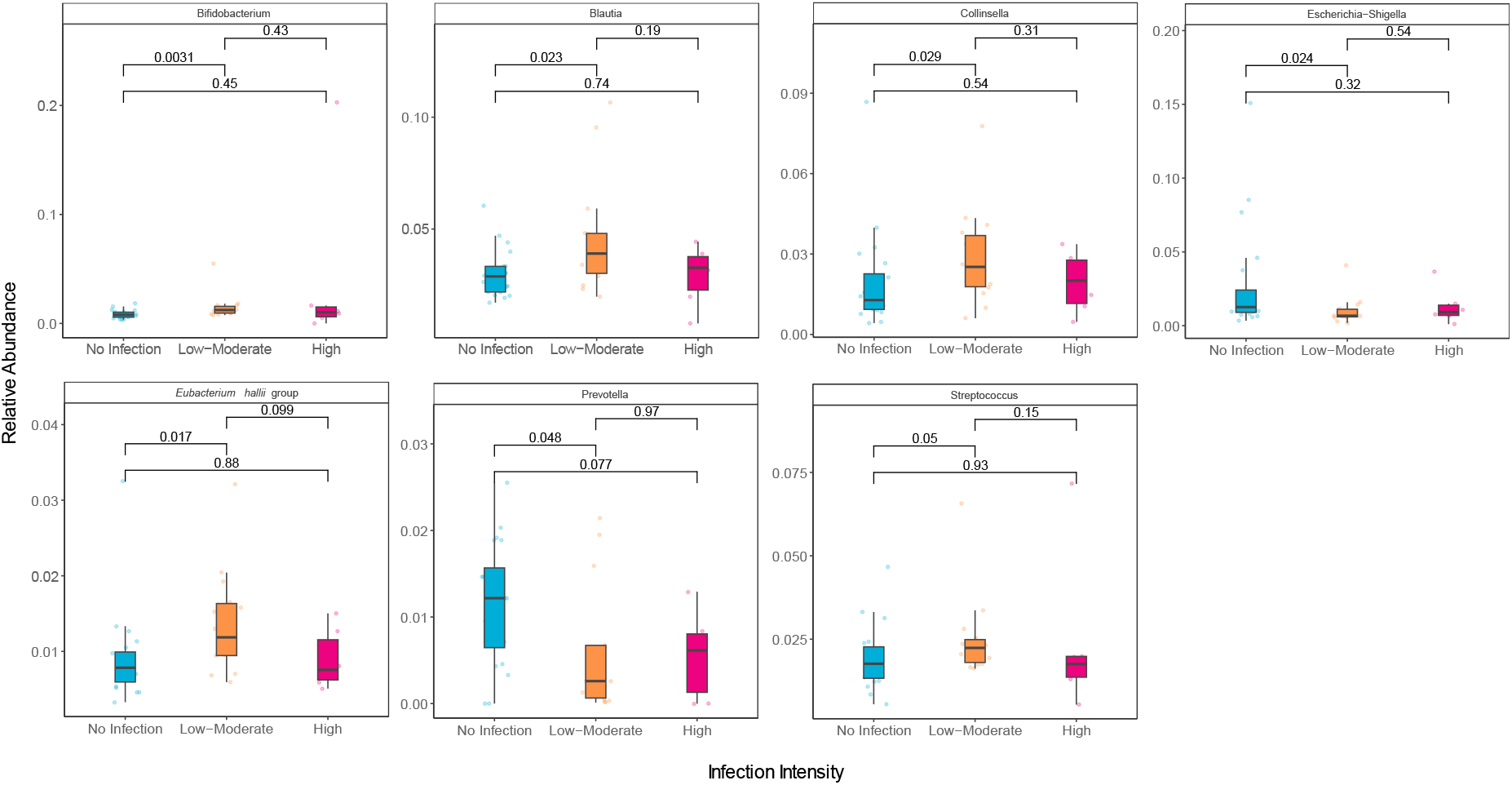
Box plots comparing the three differentially abundant microbiota within samples grouped according to infection burden. Negative (blue), Low-moderate infection intensity (orange) and High infection intensity (red).

We further investigated to determine whether these compositional shifts in the microbiome were also apparent in accordance with infection intensity. Analysis based on infection intensity revealed an increased abundance of the two main genera of Actinomycetota: *Bifidobacterium* and *Collinsella*. In low-moderate *S. mansoni* infection samples, both *Bifidobacterium* (*p*= 0.003) and *Collinsella* (*p*= 0.029) had higher relative abundances when compared with uninfected samples. In addition to this, taxa *Eubacterium hallii* (*p*= 0.017), *Blautia* (*p*= 0.023), and *Streptococcus* (*p*= 0.05) also had higher relative abundances among infected when compared with uninfected samples. On the other hand, *Prevotella* (*p*= 0.048) and *Escherichia-Shigella* (*p*= 0.024) were also significantly reduced in low-moderate *S. mansoni* infection samples. Notably, none of the observed alterations in the relative abundances of these taxa were significantly associated with the highly infected samples (>400 EPG) (p>0.05). (Fig. 3).

## Discussion

In this study, we concentrated on corroborating the association between changes in the composition and diversity of the gut microbiome, *S. mansoni* infection, and *S. mansoni* infection burden. A growing body of evidence shows that helminth infections result in dysbiosis of the gut microbiome [5,20–23]. However, the changes in the composition of individual microbiota remain highly variable [21–23]. There is a lack of agreement on the effect of *S. mansoni* infection on microbiota diversity of the gut microbiome. While some studies have shown that *S. mansoni* infection causes a reduction in alpha diversity [1,24], others have reported an increase in the alpha diversity of the gut microbiome in *S. mansoni*-infected individuals [25] or failed to find a consistent association altogether. In our study, there were no differences in alpha diversity between *S. mansoni-*infected and uninfected individuals. The underlying discrepancies leading to the varying outcomes in the alpha diversity of *S. mansoni-*infected people remain unclear, which underscores the need for further research. Nonetheless, we believe that differences in demographic characteristics of the sampled populations, sample sizes, and methodological approaches could potentially contribute to disparities in reported alpha diversities [26].

Microbiota of the genera *Bifidobacterium* and *Collinsella* were significantly elevated in *S. mansoni-* infected samples with low to moderate infection intensities compared to the negatives. *Bifidobacterium* species are known probiotics [27] and have been shown to improve overall gut health through their immune regulatory activity and ability to regulate and enhance epithelial cell turnover by the production of short-chain fatty acids (SCFA) and high concentrations of acetate [28]. Probiotics have shown great utility in the control of helminth zoonosis. The probiotic *Lactobacillus sporogenes* has been shown to decrease *S. mansoni*-induced DNA damage in mice [29]. A combination of *Streptococcus thermophilus, Lactobacillus acidophilus*, and *Bifidobacterium bifidum*, when administered to *S. mansoni* infected mice, led to a 50%–66% decrease in worm load along with a 70% and 56% reduction in *S. mansoni* egg count in liver and intestine, respectively [30].

The abundance of *Bifidobacterium* in low-moderate infections is most likely due to a host adaptation, which is linked to its probiotic activity. During *S. mansoni* infection, soluble egg antigens stimulate the production of proinflammatory cytokines [31]. *Bifidobacterium* species play a crucial role in anti-inflammatory processes by reducing IL-6 and Monocyte Chemoattractant Protein-1 (MCP-1) [32], which controls the movement of monocytes/ macrophages. They also influence the Th-2 cell immune response, boost regulatory T-cell activity [33], and enhance the availability of eosinophils and IgE antibodies [34]. These mechanisms counteract inflammation, thereby offering protection against various diseases, including inflammatory bowel diseases (Crohn’s disease, ulcerative colitis), enteric infections, colorectal cancer, diarrhoea and depression.[35–37].

In *Schistosoma mansoni* infection, both juvenile and adult worms utilise host-derived glucose and stored glycogen to fuel their growth and maturation processes [38]. This metabolic adaptation involves the downregulation of key rate-limiting enzymes, notably glycogen synthase, resulting in enhanced depletion of glycogen reserves and consequent elevation of glucose bioavailability [39]. Although the mechanism of action remains unknown, evidence suggests that *Collinsella* disrupts host glycogenic pathways while modulating other metabolic processes [40]. As such, the increased abundance of *Collinsella* in low-to-moderately infected individuals in our study may indicate a possible role of *Collinsella* in facilitating parasite growth dynamics.

Interestingly, the pathobiont *Escherichia-Shigella* was significantly reduced in infected samples. *Bifidobacterium* strains have been shown to confer host protection from enteropathogenic bacterial infections caused by *Escherichia coli* and *Shigella* through the production of high concentrations of acetate [41]. We reason that the high abundance of *Bifidobacterium* in infected samples could be directly suppressing the proliferation of the pathobiont.

A notable finding of our study was the selective increase of certain key microbiota in individuals with low-moderate *S. mansoni* infection, which was not observed in those with high infection burdens. The helminth infection load likely plays a crucial role in the intricate interactions between the host and the microbiome [42]. A heavy helminth burden in the gut may overstimulate the host’s immune system, triggering excessive inflammation and disrupting immune homeostasis [43]. This inflammatory dysregulation could compromise gut barrier function, significantly alter nutrient availability, or affect the mucus layer, all of which are critical for supporting the growth of beneficial bacteria, such as *Clostridia*, which thrive in a low-inflammatory environment [44].

In contrast, in individuals with low-moderate infections, the immune system may be only mildly activated, promoting a more tolerogenic immune environment [45]. This balanced immune response could facilitate the proliferation of beneficial bacteria. Additionally, factors such as competition for nutrients [46] and helminth load-dependent mechanical alterations to the gut, including localised inflammation, tissue damage, or structural remodelling [47] may further contribute to the distinct microbiota patterns observed across different infection intensities. However, further research is required to definitively show this, as our sample size, particularly of the high-intensity burden infections, may prove to be a limitation of the study.

## Conclusion

Our findings offer insight into the differential changes within the human gut microbiome during *Schistosoma* infections and how parasite loads are associated with the enrichment of different microbiota. The information gained from this study could be explored further to develop an alternate approach or adjunct therapy to treat intestinal schistosomiasis.

## Acknowledgements

We appreciate the NIINE laboratory and field work teams, the Council for Scientific and Industrial Research faculty, community leaders and participants in Nyive.

## Notes

### Competing Interest Statement

The authors have declared no competing interest.

## References

1. Jenkins TP, Peachey LE, Ajami NJ, MacDonald AS, Hsieh MH, Brindley PJ, et al. Schistosoma mansoni infection is associated with quantitative and qualitative modifications of the mammalian intestinal microbiota. Sci Rep. 2018;8. doi:10.1038/s41598-018-30412-x

2. Hailu T, Alemu M, Abera B, Mulu W, Yizengaw E, Genanew A, et al. Multivariate analysis of factors associated with Schistosoma mansoni and hookworm infection among primary school children in rural Bahir Dar, Northwest Ethiopia. Trop Dis Travel Med Vaccines. 2018;4. doi:10.1186/s40794-018-0064-6

3. Rujeni N, Morona D, Ruberanziza E, Mazigo HD. Schistosomiasis and soil-transmitted helminthiasis in Rwanda: An update on their epidemiology and control. Infectious Diseases of Poverty. BioMed Central Ltd.; 2017. doi:10.1186/s40249-016-0212-z

4. Holzscheiter M, Layland LE, Loffredo-Verde E, Mair K, Vogelmann R, Langer R, et al. Lack of host gut microbiota alters immune responses and intestinal granuloma formation during schistosomiasis. Clin Exp Immunol. 2014;175:246–257. doi:10.1111/cei.12230

5. Martin I, Kaisar MMM, Wiria AE, Hamid F, Djuardi Y, Sartono E, et al. The Effect of Gut Microbiome Composition on Human Immune Responses: An Exploration of Interference by Helminth Infections. Front Genet. 2019;10. doi:10.3389/fgene.2019.01028

6. Zhao Y, Yang S, Li B, Li W, Wang J, Chen Z, et al. Alterations of the mice gut microbiome via schistosoma japonicum ova-induced granuloma. Front Microbiol. 2019;10. doi:10.3389/fmicb.2019.00352

7. Costain AH, MacDonald AS, Smits HH. Schistosome Egg Migration: Mechanisms, Pathogenesis and Host Immune Responses. Frontiers in Immunology. Frontiers Media S.A.; 2018. doi:10.3389/fimmu.2018.03042

8. Jiao Y, Wu L, Huntington ND, Zhang X. Crosstalk Between Gut Microbiota and Innate Immunity and Its Implication in Autoimmune Diseases. Frontiers in Immunology. Frontiers Media S.A.; 2020. doi:10.3389/fimmu.2020.00282

9. Wiertsema SP, van Bergenhenegouwen J, Garssen J, Knippels LMJ. The interplay between the gut microbiome and the immune system in the context of infectious diseases throughout life and the role of nutrition in optimizing treatment strategies. Nutrients. MDPI AG; 2021. pp. 1– 14. doi:10.3390/nu13030886

10. Schneeberger PHH, Gueuning M, Welsche S, Hürlimann E, Dommann J, Häberli C, et al. Different gut microbial communities correlate with efficacy of albendazole-ivermectin against soil-transmitted helminthiases. Nature Communications 2022 13:1. 2022;13: 1–12. doi:10.1038/s41467-022-28658-1

11. Stark KA, Rinaldi G, Cortés A, Costain A, MacDonald AS, Cantacessi C. The role of the host gut microbiome in the pathophysiology of schistosomiasis. Parasite Immunology. John Wiley and Sons Inc; 2023. doi:10.1111/pim.12970

12. Magalhães RJS, Biritwum NK, Gyapong JO, Brooker S, Zhang Y, Blair L, et al. Mapping Helminth co-infection and co-intensity: Geostatistical prediction in Ghana. PLoS Negl Trop Dis. 2011;5. doi:10.1371/journal.pntd.0001200

13. Illumina. 16S Metagenomic Sequencing Library Preparation. 2013 [cited 14 May 2022]. Available: https://support.illumina.com/documents/documentation/chemistry_documentation/16s/16s-metagenomic-library-prep-guide-15044223-b.pdf

14. Bolyen E, Rideout JR, Dillon MR, Bokulich NA, Abnet CC, Al-Ghalith GA, et al. Reproducible, interactive, scalable and extensible microbiome data science using QIIME 2. Nature Biotechnology 2019 37:8. 2019;37:852–857. doi:10.1038/s41587-019-0209-9

15. Ssekagiri A, Sloan W, Ijaz U. microbiomeSeq: An R package for analysis of microbial communities in an environmental context. 2017. doi:10.13140/RG.2.2.17108.71047

16. Chao A, Gotelli NJ, Hsieh TC, Sander EL, Ma KH, Colwell RK, et al. Rarefaction and extrapolation with Hill numbers: a framework for sampling and estimation in species diversity studies. Ecol Monogr. 2014. Available: http://purl.oclc.org/estimates

17. Dixon P. VEGAN, a package of R functions for community ecology. Journal of Vegetation Science. 2003;14:927–930. doi:10.1111/j.1654-1103.2003.tb02228.x

18. R Core Team. R: A Language and Environment for Statistical Computing. 2013. Available: http://www.R-project.org/

19. Shuangbin Xu, Guangchuang Yu. MicrobiotaProcess: A comprehensive R package for managing and analyzing microbiome and other ecological data within the tidy framework. In: R package version 1.8.1 [Internet]. 2022. Available: https://github.com/YuLab-SMU/MicrobiotaProcess/

20. Jenkins TP, Rathnayaka Y, Perera PK, Peachey LE, Nolan MJ, Krause L, et al. Infections by human gastrointestinal helminths are associated with changes in faecal microbiota diversity and composition. PLoS One. 2017;12. doi:10.1371/journal.pone.0184719

21. Amaruddin AI, Hamid F, Koopman JPR, Muhammad M, Brienen EAT, van Lieshout L, et al. The bacterial gut microbiota of schoolchildren from high and low socioeconomic status: A study in an urban area of makassar, indonesia. Microorganisms. 2020;8:1–12. doi:10.3390/microorganisms8060961

22. Easton A V., Quiñones M, Vujkovic-Cvijin I, Oliveira RG, Kepha S, Odiere MR, et al. The impact of anthelmintic treatment on human gut microbiota based on cross-sectional and pre- and postdeworming comparisons in Western Kenya. mBio. 2019;10:1–14. doi:10.1128/MBIO.00519-19

23. Huwe T, Prusty BK, Ray A, Lee S, Ravindran B, Michael E. Interactions between parasitic infections and the human gut microbiome in Odisha, India. American Journal of Tropical Medicine and Hygiene. 2019;100:1486–1489. doi:10.4269/ajtmh.18-0968

24. Schneeberger PHH, Coulibaly JT, Panic G, Daubenberger C, Gueuning M, Frey JE, et al. Investigations on the interplays between Schistosoma mansoni, praziquantel and the gut microbiome. Parasit Vectors. 2018;11. doi:10.1186/s13071-018-2739-2

25. Ajibola O, Rowan AD, Ogedengbe CO, Mshelia MB, Cabral DJ, Eze AA, et al. Urogenital schistosomiasis is associated with signatures of microbiome dysbiosis in Nigerian adolescents. Sci Rep. 2019;9. doi:10.1038/s41598-018-36709-1

26. Liu K, Zhang Y, Li Q, Li H, Long D, Yan S, et al. Ethnic differences shape the Alpha but not beta diversity of gut microbiota from school children in the absence of environmental differences. Microorganisms. 2020;8. doi:10.3390/microorganisms8020254

27. Ray D, Alpini G, Glaser S. Probiotic Bifidobacterium species: potential beneficial effects in diarrheal disorders. Focus on “Probiotic Bifidobacterium species stimulate human SLC26A3 gene function and expression in intestinal epithelial cells.” Am J Physiol Cell Physiol. 2014;307:C1081. doi:10.1152/AJPCELL.00300.2014

28. Binda C, Lopetuso LR, Rizzatti G, Gibiino G, Cennamo V, Gasbarrini A. Actinobacteria: A relevant minority for the maintenance of gut homeostasis. Dig Liver Dis. 2018;50:421–428. doi:10.1016/J.DLD.2018.02.012

29. Mohamed AH, Osman GY, Zowail MEM, El-Esawy HMI. Effect of Lactobacillus sporogenes (probiotic) on certain parasitological and molecular aspects in Schistosoma mansoni infected mice. J Parasit Dis. 2016;40:823. doi:10.1007/S12639-014-0586-4

30. Abdel-Salam AM, Ammar N, Abdel-Hamid AZ. Effectiveness of Probiotic Labneh Supplemented with Garlic or Onion Oil Against Schistosoma mansoni in Infected Mice. International Journal of Dairy Science. 2008;3:97–104. doi:10.3923/ijds.2008.97.104

31. Masamba P, Kappo AP. Immunological and Biochemical Interplay between Cytokines, Oxidative Stress and Schistosomiasis. International Journal of Molecular Sciences 2021, Vol 22, Page 7216. 2021;22:7216. doi:10.3390/IJMS22137216

32. Moya-Pérez A, Neef A, Sanz Y. Bifidobacterium pseudocatenulatum CECT 7765 Reduces Obesity-Associated Inflammation by Restoring the Lymphocyte-Macrophage Balance and Gut Microbiota Structure in High-Fat Diet-Fed Mice. PLoS One. 2015;10:e0126976. doi:10.1371/JOURNAL.PONE.0126976

33. Henrick BM, Rodriguez L, Lakshmikanth T, Pou C, Henckel E, Arzoomand A, et al. Bifidobacteria-mediated immune system imprinting early in life. Cell. 2021;184:3884-3898.e11. doi:10.1016/J.CELL.2021.05.030

34. Rosenberg HF, Masterson JC, Furuta GT. Eosinophils, probiotics, and the microbiome. J Leukoc Biol. 2016;100:881–888. doi:10.1189/JLB.3RI0416-202R

35. O’Callaghan A, van Sinderen D. Bifidobacteria and Their Role as Members of the Human Gut Microbiota. Front Microbiol. 2016;7:925. doi:10.3389/FMICB.2016.00925

36. Arboleya S, Watkins C, Stanton C, Ross RP. Gut bifidobacteria populations in human health and aging. Front Microbiol. 2016;7:1204. doi:10.3389/FMICB.2016.01204/BIBTEX

37. Cheng LH, Liu YW, Wu CC, Wang S, Tsai YC. Psychobiotics in mental health, neurodegenerative and neurodevelopmental disorders. J Food Drug Anal. 2019;27:632. doi:10.1016/J.JFDA.2019.01.002

38. You H, Stephenson RJ, Gobert GN, McManus DP. Revisiting glucose uptake and metabolism in schistosomes: New molecular insights for improved schistosomiasis therapies. Frontiers in Genetics. Frontiers Research Foundation; 2014. doi:10.3389/fgene.2014.00176

39. von Bülow V, Gindner S, Baier A, Hehr L, Buss N, Russ L, et al. Metabolic reprogramming of hepatocytes by Schistosoma mansoni eggs. JHEP Reports. 2023;5. doi:10.1016/j.jhepr.2022.100625

40. Gomez-Arango LF, Barrett HL, Wilkinson SA, Callaway LK, McIntyre HD, Morrison M, et al. Low dietary fiber intake increases Collinsella abundance in the gut microbiota of overweight and obese pregnant women. Gut Microbes. 2018;9:189–201. doi:10.1080/19490976.2017.1406584

41. Fukuda S, Toh H, Taylor TD, Ohno H, Hattori M. Acetate-producing bifidobacteria protect the host from enteropathogenic infection via carbohydrate transporters. https://doi.org/104161/gmic21214. 2012;3:p449–454. doi:10.4161/GMIC.21214

42. Zaiss MM, Harris NL. Interactions between the intestinal microbiome and helminth parasites. Parasite Immunol. 2016;38:5–11. doi:10.1111/pim.12274

43. McSorley HJ, Maizels RM. Helminth infections and host immune regulation. Clin Microbiol Rev. 2012;25:585–608. doi:10.1128/CMR.05040-11

44. Vacca F, Le Gros G. Tissue-specific immunity in helminth infections. Mucosal Immunology. Springer Nature; 2022. pp. 1212–1223. doi:10.1038/s41385-022-00531-w

45. Gazzinelli-Guimaraes PH, Nutman TB. Helminth parasites and immune regulation. F1000Research. F1000 Research Ltd; 2018. doi:10.12688/F1000RESEARCH.15596.1

46. Horrocks V, King OG, Yip AYG, Marques IM, McDonald JAK. Role of the gut microbiota in nutrient competition and protection against intestinal pathogen colonization. Microbiology (United Kingdom). Microbiology Society; 2023. doi:10.1099/mic.0.001377

47. Peachey LE, Jenkins TP, Cantacessi C. This Gut Ain’t Big Enough for Both of Us. Or Is It? Helminth–Microbiota Interactions in Veterinary Species. Trends in Parasitology. Elsevier Ltd; 2017. pp. 619–632. doi:10.1016/j.pt.2017.04.004

